# Functional diversity of GPCR-Gustducin complexes controls signaling output and suppresses alternative pathways

**DOI:** 10.64898/2026.03.28.715036

**Authors:** Victoria Jaime Arce, Mona Stefanie Tögl, Santiago Javier Bonn Garcia, Isabell Kaczmarek, Daniel Hilger, Hannes Schihada

**Author notes:** contributed equally (Victoria Jaime Arce, Mona Stefanie Tögl).

## Abstract

Gustducin (G_gust_), the G protein mediating taste receptor signaling, is expressed not only in sensory taste cells but throughout the human body alongside numerous non-taste G protein-coupled receptors (GPCRs). Whether and how these receptors engage G_gust_, and the functional consequences of such interactions, remain poorly understood. Here, we developed two complementary biosensors that enable direct monitoring of GPCR-G_gust_ interaction and downstream signaling in living cells, allowing pharmacological characterization of G_gust_ activity by non-taste GPCRs. Unexpectedly, we find that several non-taste GPCRs do not activate G_gust_ but instead stabilize the G protein heterotrimer and suppress basal signaling by forming unproductive GPCR-G protein complexes, whereas others induce conventional Gα_gust_ activation and signal transduction. We further show that these unproductive GPCR-G_gust_ complexes suppress the activation of alternative signaling pathways by sequestering receptors from other G proteins. This previously unrecognized mode of interaction functionally partitions GPCRs into activators and inactivators of G_gust_ and represents a mechanism for balancing concurrent signaling pathways within a cell.

**Highlights:**

- Development of biosensors for gustducin activation and signal transduction
- Analysis of single-cell co-expression data of non-tastant GPCRs and gustducin
- Profiling of non-tastant GPCR-induced gustducin activation and signal transduction
- Productive and unproductive GPCR-gustducin complexes revealed
- Unproductive GPCR-gustducin complexes suppress alternative signaling pathways

## Introduction

Gα_gustducin_ (Gα_gust_), encoded by *GNAT3*, is a member of the G_i/o_ family of G proteins. Gα_gust_ is the major component of the G protein heterotrimer G_gust_ (Gα_gust_βγ), which is composed of Gα_gust_ and a Gβγ dimer, and plays a central role in taste perception by coupling bitter (TAS2Rs) and sweet/umami (TAS1Rs) tastant-binding G protein-coupled receptors (GPCRs) to intracellular signaling pathways[1–3]. Gα_gust_ is closely related to the transducins Gα_t1_ and Gα_t2_ (encoded by *GNAT1* and *GNAT2*, respectively), which mediate vision in rod and cone photoreceptor cells. Transducins and gustducin, as well as other G_i/o_-family Gα subunits and Gα_s_, can enhance phosphodiesterase (PDE) activity by binding mainly to the inhibitory γ-subunit of PDEs[4–7]. This interaction relieves autoinhibition of the catalytic subunits, resulting in increased PDE activity and, consequently, reduced intracellular levels of the cyclic nucleotides cAMP and cGMP. In addition, Gβγ released from activated Gα_gust_ activates phospholipase Cβ2 (PLCβ2), promoting IP_3_- and Ca^2+^-dependent signaling and opening of transient receptor potential cation channels subfamily M member 5 (TRPM5) channels, which leads to depolarization and neurotransmitter release[8–10]. In addition to its role in type II taste receptor cells of the tongue and palate, Gα_gust_ is expressed in specialized epithelial cells throughout the stomach, small intestine, and colon. Here, Gα_gust_ couples taste receptors to glucagon-like peptide 1 (GLP-1) secretion, ghrelin release and expression of sodium-dependent glucose transporter isoform 1 (SGLT1)[11–13].

With the emergence of single-cell and single-nucleus sequencing techniques and organism-wide protein expression profiling, it is apparent that Gα_gust_ can also be co-expressed with non-taste receptors in the human body. Beyond the tongue, the human protein atlas reports Gα_gust_ expression in the brain, testis and adipose tissue, and – at high levels – in the duodenum and other parts of the small intestine[14,15]. These tissues also express numerous non-taste GPCRs. However, whether G_gust_ can couple to and mediate signaling of these receptors remains unclear.

In the present study, we addressed this knowledge gap using an integrative approach that combines the analysis of single-cell expression data with newly developed biosensors for pathway-unbiased G_gust_ activation and Gα_gust_-mediated signal transduction. Using these tools, we find that several non-taste GPCRs can couple to Gα_gust_-containing heterotrimers. Interestingly, this coupling results in two functionally distinct complexes: conventional signaling complexes that promote G_gust_ activation, such as those formed with the A_1_ adenosine receptor (A_1_R) and histamine H_3_ receptor (H_3_R), and complexes that suppress basal Gα_gust_ signal transduction, such as those formed with the histamine H_2_ receptor (H_2_R) or the A_2A_ adenosine receptor (A_2A_R). Nucleotide-sensitivity experiments confirmed these GPCR-G_gust_ interactions. These experiments suggest a previously unrecognized mechanism by which G_gust_ may modulate gastrointestinal physiological functions in response to non-taste receptor ligands.

## Results

### A new biosensor for G_gust_ heterotrimer dissociation in cells

Given the divergent signaling cascades promoted upon G_gustducin_ dissociation into Gα_gust_ and Gβγ, we first aimed to develop a signaling-pathway independent readout of G_gust_ activation. In line with the placement of energy partners in a TRUPATH G_gust_ heterotrimer dissociation sensor[16], we fused NanoLuciferase[17] (Nluc), flanked by flexible linkers, to Asp116 in Gα_gust_. Next, we combined the sequence of Gα_gust_-Nluc with the wildtype Gβ_3_ gene and N-terminally labeled Gγ_9_ on a plasmid backbone comprising a T2A self-cleavage site (between Gβ_3_ and cpVenus-Gγ_9_) and an internal ribosome entry site (IRES; upstream of Gα_gust_-Nluc), because this Gβγ dimer proved to be a sensitive module to detect the dissociation of other G protein heterotrimers[18,19]. This plasmid design enabled the simultaneous co-transfection of all three G protein subunits (**Fig. 1A, Fig. S1A**). To facilitate the future combination of this new sensor, which we refer to as Ggust-CASE, with other optical biosensors, we created a version of G-CASE with a dimmed BRET acceptor (“dark Ggust-CASE”) by mutating Tyr218 in cpVenus to Trp[20]. The quenched emission of cpVenus was confirmed in the luminescence emission spectra of transfected cells (**Fig. 1B**). Starting from dark Ggust-CASE, we also generated a Ggust-CASE variant with quenched emission of Nluc by mutating Arg164 to Gln in the luciferase. This new construct, called “double dark Ggust-CASE”, exhibited only 0.2 % of the luminescence emission compared to dark Ggust-CASE (comparison of area under the curves from 400 – 600 nm; **Fig. S2**). Next, we tested whether Ggust-CASE responds to agonist-mediated stimulation of a prototypical G_i/o_-coupled GPCR. Stimulation of the co-expressed histamine H_3_ receptor (H_3_R) with histamine resulted in a time- and concentration-dependent decrease in BRET of Ggust-CASE (**Fig. 1C, D**). The potency of histamine (mean ± SEM pEC_50_ = 7.24 ± 0.23) was consistent with the activation of other G_i/o_ biosensors by histamine/H_3_R[18].

**Figure 1.**
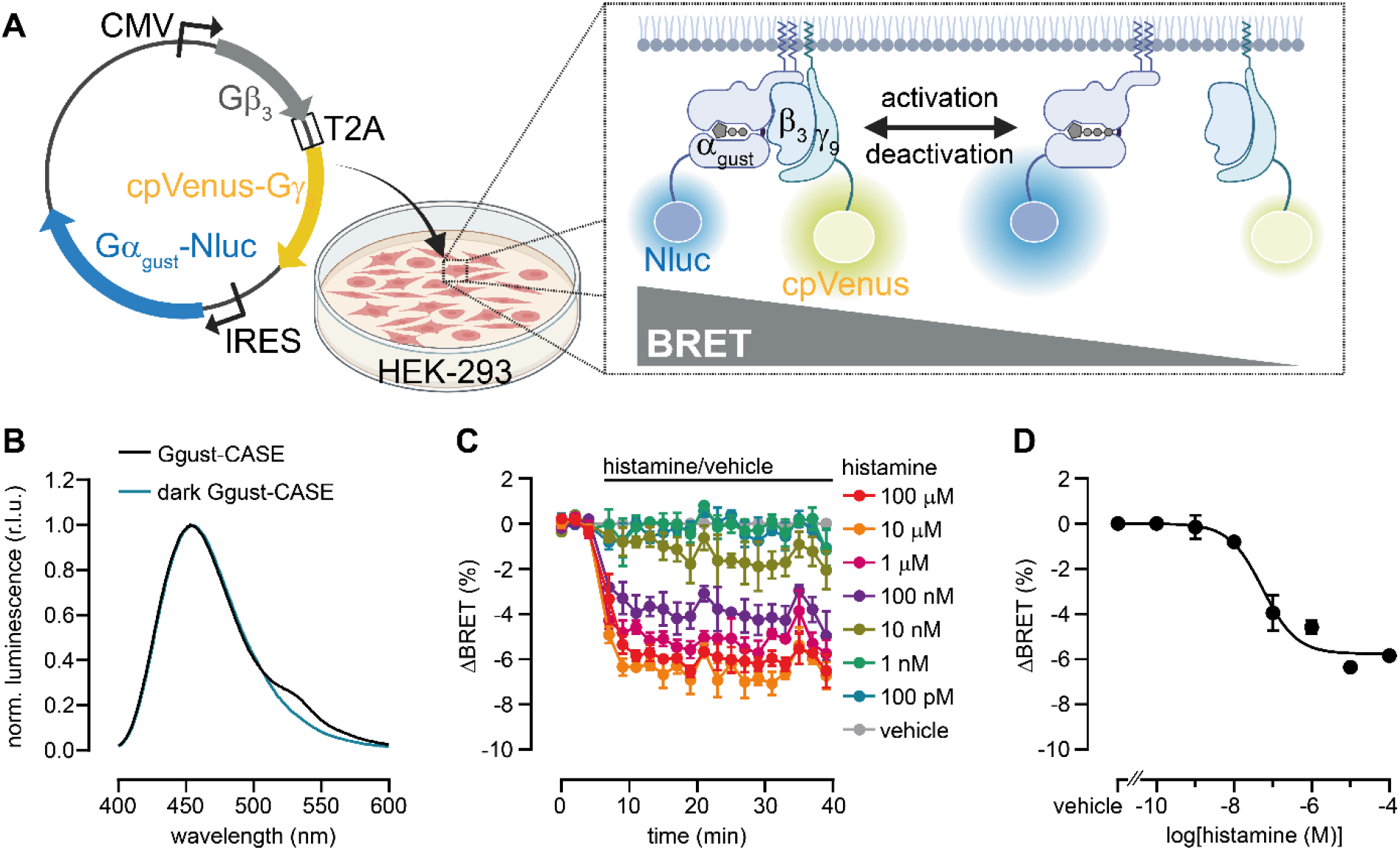
Development of the Ggust dissociation biosensor, Ggust-CASE. **A)** Scheme of the plasmid design and biosensor principle. **B)** Peak-normalized luminescence emission spectra of Ggust-CASE and dark Ggust-CASE harboring the Y218W point mutation in cpVenus. **C)** ΔBRET time course of Ggust-CASE upon stimulation of co-expressed H_3_R with histamine. **D)** ΔBRET concentration-response curve of histamine measured with Ggust-CASE in H_3_R-expressing cells. Data in B show one representative experiment. Data in C and D represent the mean ± SEM of three biological replicates. Experiments were conducted in HEK-293A cells that were transiently transfected with either Ggust-CASE (B-D) or dark Ggust-CASE (B), either without (B) or with H_3_R (C, D).

### A biosensor for Gα_gust_-mediated signal transduction in cells

To expand our G_gustducin_-targeted toolbox to include a cell-based readout of Gα_gust_-mediated signal transduction, we tested whether known effectors of the different Gα subtypes would interact with Gα_gust_ following heterotrimer dissociation. We used an intermolecular BRET setup to combine cpVenus- or YFP-labeled Gα effectors with Gα-Nluc. Our analysis included the Gα_q/11_ effectors PLCβ3, p63RhoGEF and GRK2[21–24], the Gα_12/13_ effector PDZ-RhoGEF[21] and the Gα_i/o_- and Gα_s_-effector adenylyl cyclase 5 (AC5)[25], labeled either N-terminally at the C1 domain (coupling site for Gα_i/o_) or C-terminally at the C2 domain (coupling site for Gα_s_), respectively (**Fig. S3A**). We detected GPCR agonist-induced increases in BRET for all effectors with their cognate Gα-Nluc partners. However, Gα_gust_-Nluc only exhibited a modest, histamine concentration-dependent increase in BRET with C1 domain-labeled AC5 (**Fig. S3B-G**).

Due to the low signal-to-noise ratio obtained with Gα_gust_-Nluc and YFP-AC5, we searched the literature for other potential interaction partners of active Gα_gust_. Co-immunoprecipitation experiments revealed that, similar to α- and β-transducin, Gα_gust_(GTP) physically interacts with the γ-subunit of PDE6[5–7]. We therefore co-expressed Gα_gust_-Nluc or Gα_q_-Nluc, the latter serving as a negative control, with cpVenus-labeled PDE6γ and H_3_R or the G_q_-coupled H_1_R, respectively (**Fig. 2A**). Here, histamine induced a time- and concentration-dependent increase in BRET with Gα_gust_-Nluc, but not with Gα_q_-Nluc. This is indicative of an enhanced interaction between Gα_gust_ and PDE6γ, and Gα_gust_-mediated signal transduction upon G protein activation (**Fig. 2B-D**). Histamine potency (mean ± SEM pEC_50_ = 7.92 ± 0.07) was approximately five-fold higher in the Gα_gust_–PDE6γ interaction assay than in the Ggust-CASE readout, consistent with signal amplification along the signaling cascade. We also titrated the plasmid ratios of co-transfected Gα_gust_-Nluc and cpVenus-labeled PDE6γ to identify the optimal conditions for this intermolecular BRET assay, but found no significant differences between the test conditions (**Fig. S4**).

**Figure 2.**
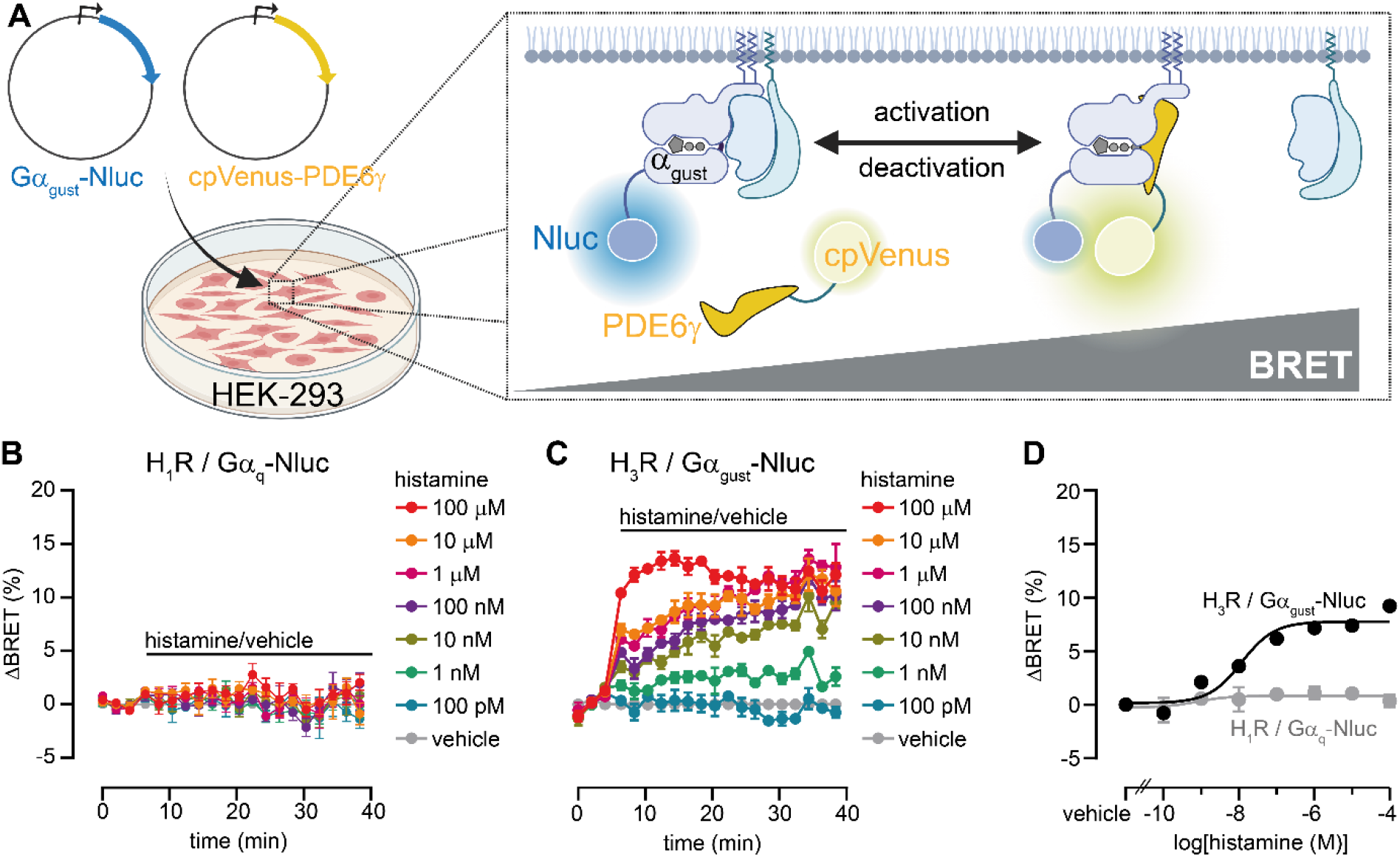
Development of a Gα_gust_-PDE6γ interaction assay. **A)** Scheme of the BRET-based protein-protein interaction assay. **B-D)** Time courses (B and C) and concentration-dependent BRET changes (D) induced by histamine in HEK-293A cells that were transiently transfected with PDE6γ-cpVenus, along with either H_1_R and Gα_q_-Nluc (B, D), or H_3_R and Gα_gust_-Nluc (C, D). Data represent the mean ± SEM of three biological replicates.

### GPCR-specific conformational responses in the G_gust_ heterotrimer

G_gust_’s primary physiological function is linked to the sensation of taste, but its expression in the epithelial cells that line the stomach, small intestine, and colon[14] suggests that it could also act as a transducer of other extracellular signals when coupled to a non-gustatory GPCR. We therefore examined expression datasets from the Human Protein Atlas to identify GPCRs that may be coupled to G_gust_ in the small intestine and duodenum. In total, 202 GPCRs are co-expressed with Gα_gust_ (*GNAT3*) in the duodenum and other parts of the small intestine (**Supplementary Data 1**). From this set of receptors, we selected 24 pharmacologically accessible GPCRs with known agonists and G protein coupling information, according to guidetopharmacology.org and gproteindb.org[26,27]. Our selection included GPCRs with G protein coupling profiles ranging from very selective (e.g., V_1B_R, P2YR_1_) to very promiscuous (e.g., BK_2_R, H_1_R) and covered all four major Gα families[28].

Next, we performed an organism-wide co-expression analysis of these receptors with *GNAT3* at the single-cell level (**Supplementary Data 2**). Our analysis of publicly available single-cell RNA sequencing data[15] confirmed that all 24 GPCRs are co-expressed with *GNAT3* at a single-cell level in humans. However, only 14 of these are co-expressed with *GNAT3* in GIT cells or cells of accessory digestive tissue (ADT) (e.g., the salivary gland). The majority of GPCRs included in our analysis (21 out of 24) were co-expressed with *GNAT3* in brain cells; a subset of these receptors also exhibited co-expression with *GNAT3* in non-GIT/non-ADT cells outside the brain (**Fig. 3A**). We subjected all 24 receptors to our cell-based pharmacological testing system, expressing them alongside Ggust-CASE in HEK293 cells in order to measure the biosensor’s response to GPCR stimulation by agonists (**Fig. 3B**). Of the 24 receptors tested, 13 GPCRs mediated agonist concentration-dependent BRET changes in Ggust-CASE. Surprisingly, GPCR activation induced not only decreases in Ggust-CASE BRET (e.g., for the adenosine A_1_ receptor, A_1_R), but positive BRET changes were also observed, similar to earlier studies with other G protein biosensors[29–31]. The increase in BRET was most prominent for the histamine H_2_ receptor (H_2_R), a primary G_s_-coupled GPCR and important regulator of gastric acid secretion in the stomach[32] – where the H_2_R is co-expressed with Gα_gust_. We profiled the H_2_R-mediated G-CASE responses of other G protein subtypes (**Fig. S5**), including other members of the G_i/o_ protein family and newly developed and validated biosensors for G_11_ and G_o2_ heterotrimers (G11-CASE and Go2-CASE; **Fig. S6, S7**), to confirm that the positive BRET change was specific to G_gust_.

**Figure 3.**
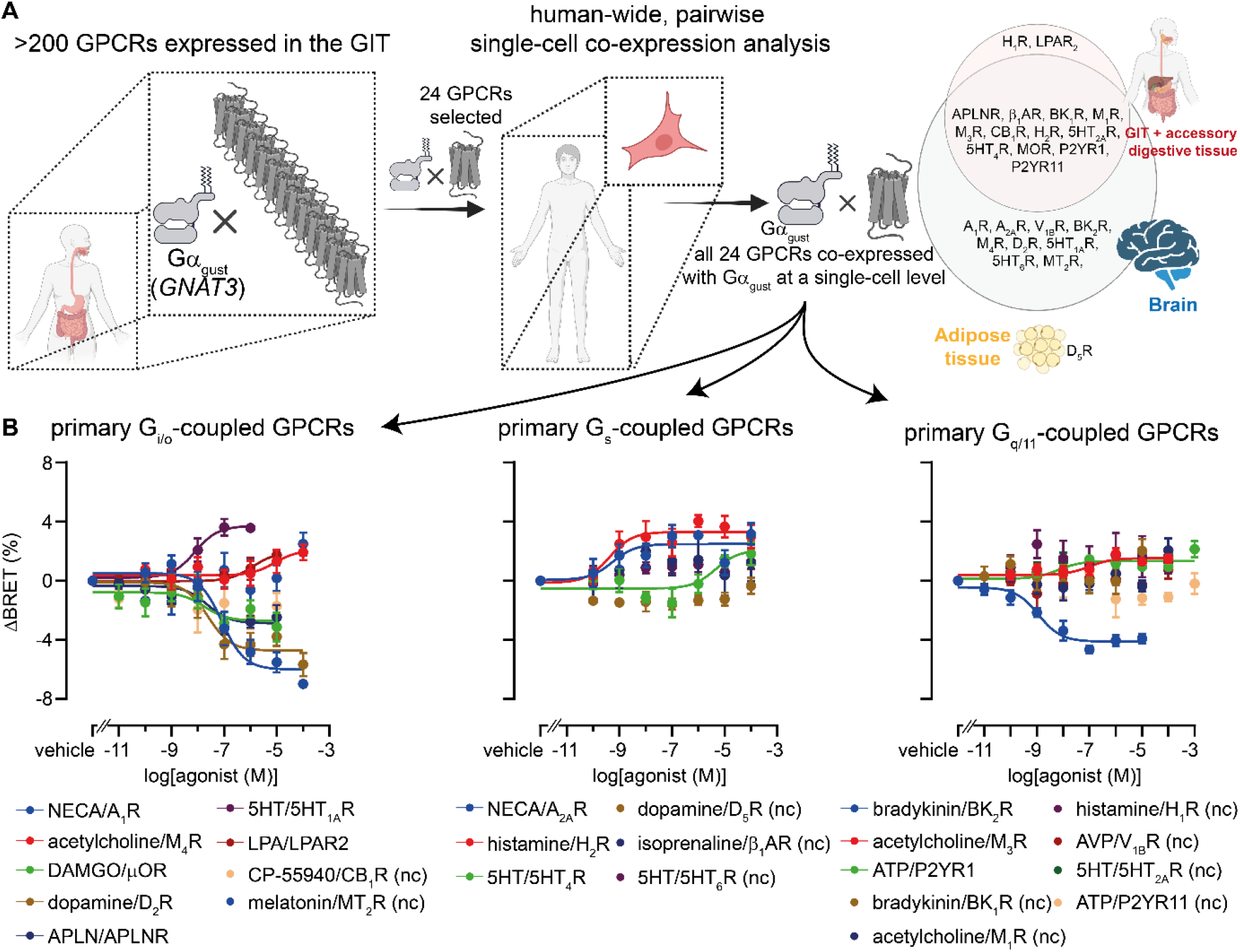
Profiling Ggust-CASE responses to gastrointestinal GPCR activation. **A)** Workflow of the co-expression analysis of GPCRs with Gα_gust_ at tissue- and single-cell levels. **B)** Concentration-dependent BRET changes induced by primary G_i/o_-, G_s_- and G_q/11_-coupled GPCRs (left to right) transiently expressed in HEK-293A cells alongside Ggust-CASE and stimulated with agonists. Data represent the mean ± SEM of three biological replicates. Non-Ggust-couplers (nc) were identified by comparing a linear model with a zero slope to a sigmoidal concentration–response model (with the Hill slope fixed to +1 or −1) using an extra sum-of-squares F-test (p > 0.05).

The receptor-dependent BRET responses of Ggust-CASE with opposing directions suggest distinct GPCR-mediated changes in conformation of the G_gust_ heterotrimer. While a decrease in FRET or BRET between Gα- and Gβγ-fused energy partners is widely accepted as representing heterotrimer dissociation and activation, an increase in FRET/BRET, as observed with the H_2_R, could be explained by receptor-mediated reduction of basal G_gust_ activity. To investigate whether Ggust exhibits basal activity in HEK-293 cells, we expressed Ggust-CASE without additional GPCRs and depleted the endogenous guanosine nucleotides GDP and GTP using apyrase following digitonin-mediated cell permeabilization[33]. After loading the G protein sensors with the apyrase-resistant GDP analog GDPβS and recording the BRET baseline of our biosensor, we added a tenfold excess of GTPγS to restore the GDP/GTP ratio present in intact cells (**Fig. 4A**). GTPγS induced a concentration-dependent decrease in Ggust-CASE BRET (**Fig. 4B, C**). The apparently low potency of GTPγS observed in our experiments (mean ± SEM pEC_50_ = 5.52 ± 0.12) compared to biochemical analyses of purified Gα proteins (e.g., k_*D*_ of GTPγS at purified Gα_t1_ 50 – 100 nM;[34]) may be explained by the presence of Gβγ and other modulators of G protein activity, such as regulators of G protein signaling (RGS) proteins and guanine nucleotide dissociation inhibitors (GDIs)[35,36], in our experimental setup. The decrease in BRET induced by GTPγS indicates that G_gust_ is at least partially active in intact cells prior to the stimulation of GPCRs by an agonist.

**Figure 4.**
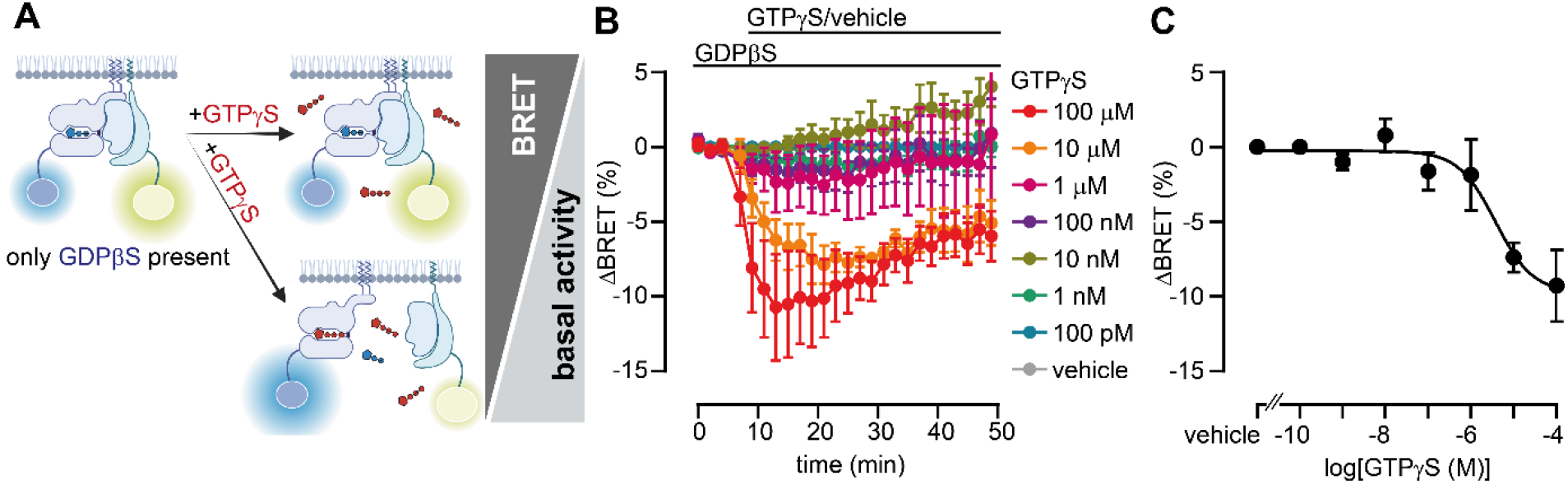
Confirmation of basal G_gust_ activity. **A)** Scheme showing the principle of the assay to determine the basal activity of G_gust_ in permeabilized and endogenous nucleotide-depleted cells. **B, C)** Time- and concentration-dependent BRET changes induced by GTPγS in HEK-293A cells that have been transiently transfected with Ggust-CASE. Data represent the mean ± SEM of three biological replicates.

To validate our hypothesis that receptors mediate the suppression of basal G_gust_ activity, we used our new Gα_gust_-PDE6γ interaction assay to probe a selected subset of receptors. This included the BK_2_R and A_1_R, which both reduced BRET in Ggust-CASE, as well as H_2_R, A_2A_R and 5HT_1A_R, which all promoted an increase in Ggust-CASE BRET. Experiments were performed using dark Ggust-CASE (carrying the loss-of-fluorescence mutation Y218W in cpVenus and wildtype Nluc on Gα_gust_) and PDE6γ-cpVenus. Interestingly, we found that the receptors that induced an increase in Ggust-CASE, reduced the interaction between Gα_gust_ and PDE6γ, and *vice versa* (**Fig. 5A**). This correlation suggests (i) basal interaction between Gα_gust_ and PDE6γ already prior to GPCR activation by an agonist and that (ii) the activation of these receptors by their agonists shifts the balance of G_gust_ activity towards inactive states. Using the experimental setup of permeabilized and nucleotide-controlled cells that we established earlier, we confirmed the basal interaction between Gα_gust_ and PDE6γ. The addition of GTPγS to GDPβS-preloaded cells enhanced BRET between these labeled interaction partners (**Fig. 5B**), suggesting that Gα_gust_ and PDE6γ already interact prior to GPCR-mediated G_gust_ activation in intact cells, where both GDP and GTP are present.

**Figure 5.**
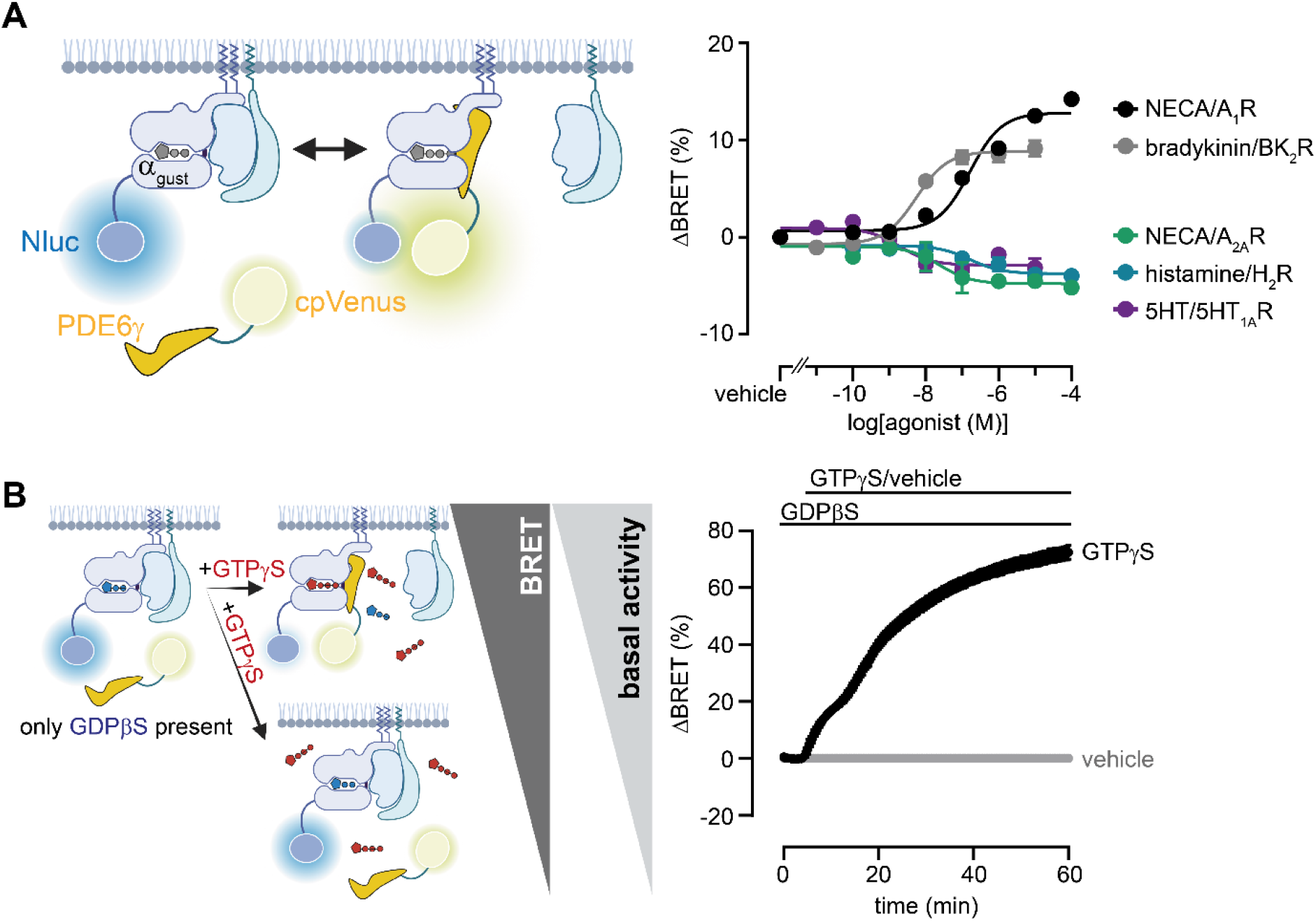
GPCR-mediated G_gust_ activation can enhance or reduce the basal interaction between Gα_gust_ and PDE6γ interaction. **A)** Scheme of the assay principle and concentration-dependent BRET changes induced by GPCR agonists in HEK-293A cells that have been transiently transfected with the indicated GPCRs, dark Ggust-CASE and PDE6γ-cpVenus. **B)** Scheme of the assay principle and GTPγS-induced BRET changes in HEK-293A cells that have been transiently transfected with dark Ggust-CASE and PDE6γ-cpVenus. Data represent the mean ± SEM of three biological replicates.

### Productive and unproductive GPCR-G_gust_ complexes revealed by nucleotide sensitivity experiments

The enhancement and reduction of the GPCR-specific Gα_gust_-PDE6γ interaction were found to be anti-correlated with Ggust-CASE BRET responses. This suggests that some GPCRs form productive complexes with G_gust_, while others form unproductive complexes, as previously described for some G_13_-coupled GPCRs[37] and virus-encoded, G_i_-coupled receptors[38]. To test this hypothesis, we adapted an established intermolecular BRET approach, which detects the proximity of a luciferase-labeled GPCR with fluorescent Gβ_1_γ_2_[37,39,40], to assay the nucleotide sensitivity of receptor-G_gust_ engagement. The C-terminally Nluc-tagged receptors were co-expressed the Venus-labeled Gβγ dimer and wildtype Gα_gust_ in HEK-293 cells lacking endogenous Gα subtypes[41]. Due to the enhanced affinity of nucleotide-free G proteins for active-state receptors[33], depleting endogenous nucleotides with apyrase and stimulating GPCRs with agonists should stabilize a high BRET state between GPCR-Nluc and fluorescently-labeled G proteins. The addition of GTPγS would then dissociate productive GPCR-G protein complexes (leading to a decrease in BRET). In contrast, the addition of GTPγS to unproductive GPCR-G_gust_ pairings, would either (i) rearrange or (ii) have no effect on preformed complexes, or (iii) enhance the assembly of these GPCR-G_gust_ complexes. All of these scenarios would lead to no change or an increase in BRET between GPCR-Nluc and Venus-Gβγ and these unproductive complexes would not be able to promote Gα(GTP)- or Gβγ-mediated signal transduction (**Fig. 6A**). We performed this analysis with the presumed productive G_gust_-couplers H_3_R, BK_2_R and A_1_R, as well as the presumed unproductive G_gust_-couplers A_2A_R, H_2_R and 5HT_1A_R. GTPγS dissociated the receptor complexes with G_gust_ for H_3_R and A_1_R, while BK_2_R exhibited a biphasic behavior, indicating initial complex rearrangement followed by dissociation (**Fig. 6B**). In contrast, enhanced complex formation or complex rearrangement was observed for A_2A_R, H_2_R and 5HT_1A_R (**Fig. 6C**). These results are consistent with our findings obtained using biosensors for G_gust_ heterotrimer dissociation (Ggust-CASE) and Gα_gust_-mediated signal transduction (Gα_gust_-PDE6γ interaction). Taken together, our pharmacological data show that G_gust_ can form both productive and unproductive complexes with GPCRs that are co-expressed in the same cell in the GIT or brain, when tested in HEK293 cells.

**Figure 6.**
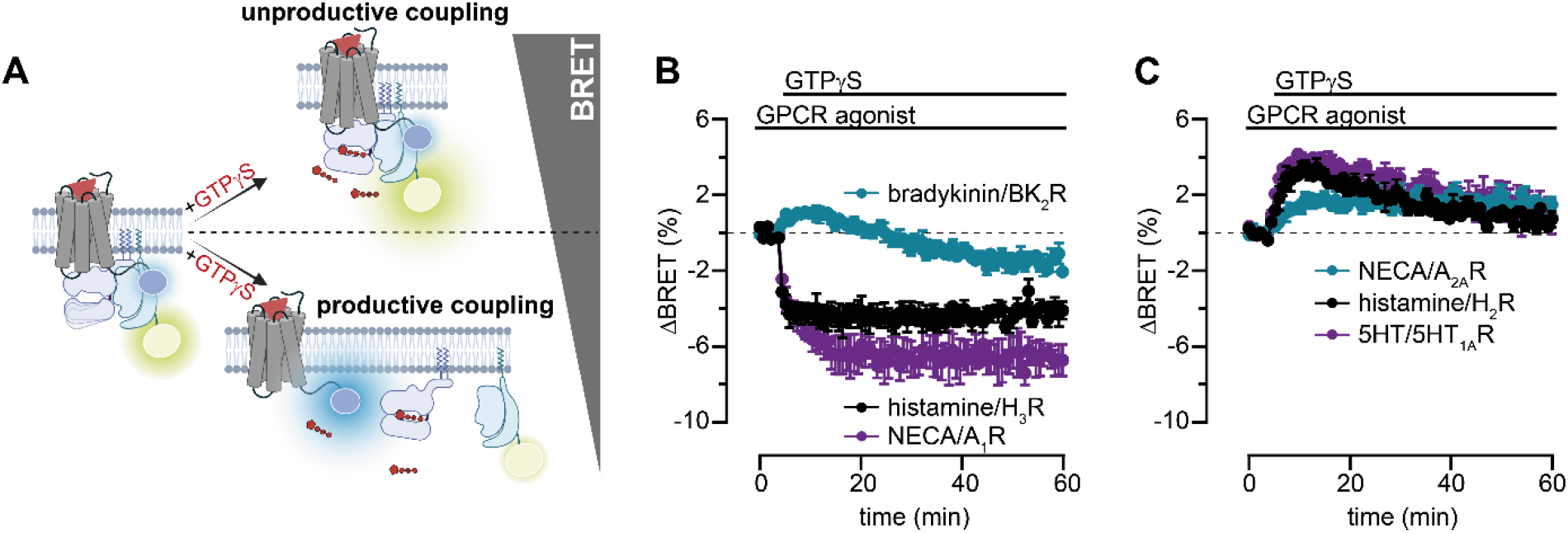
Nucleotide sensitivity of GPCR-G_gust_ complexes. **A)** Scheme of the assay principle. **B, C)** GTPγS-induced BRET changes between G_gust_ heterotrimers and supposedly productive (B) and unproductive (C) G_gust_ couplers. All experiments were conducted in endogenous Gα-deficient HEK-293A cells that had been transiently transfected with the indicated GPCR-Nluc constructs, as well as Gα_gust_ and Venus-labeled Gβ_1_γ_2_. Data represent the mean ± SEM of three biological replicates.

### Unproductive GPCR-G_gust_ complexes sequester receptors from other G proteins

The formation of unproductive GPCR-Ggust complexes may have important implications for signal transduction in cells mediated by other G proteins or GPCR coupling partners[37] Therefore, we investigated the effect of G_gust_ heterotrimer (double dark Ggust-CASE) overexpression on H_2_R- and A_1_R-mediated activation of other G proteins and recruitment of β-arrestin2. While G_q_ activation by the A_1_R was not affected by G_gust_ overexpression (**Fig. 7A**), the amplitude of G_s(long)_ activation by the H_2_R was substantially reduced in the presence of G_gust_ (**Fig. 7B**). The same pattern was observed when receptor-mediated β-arrestin2 translocation to the plasma membrane was assayed with and without G_gust_ overexpression (**Fig. S8A-C**). Importantly, the surface expression levels of the H_2_R were similar in experiments with and without double dark Ggust-CASE co-expression (**Fig. S8D**).

**Figure 7.**
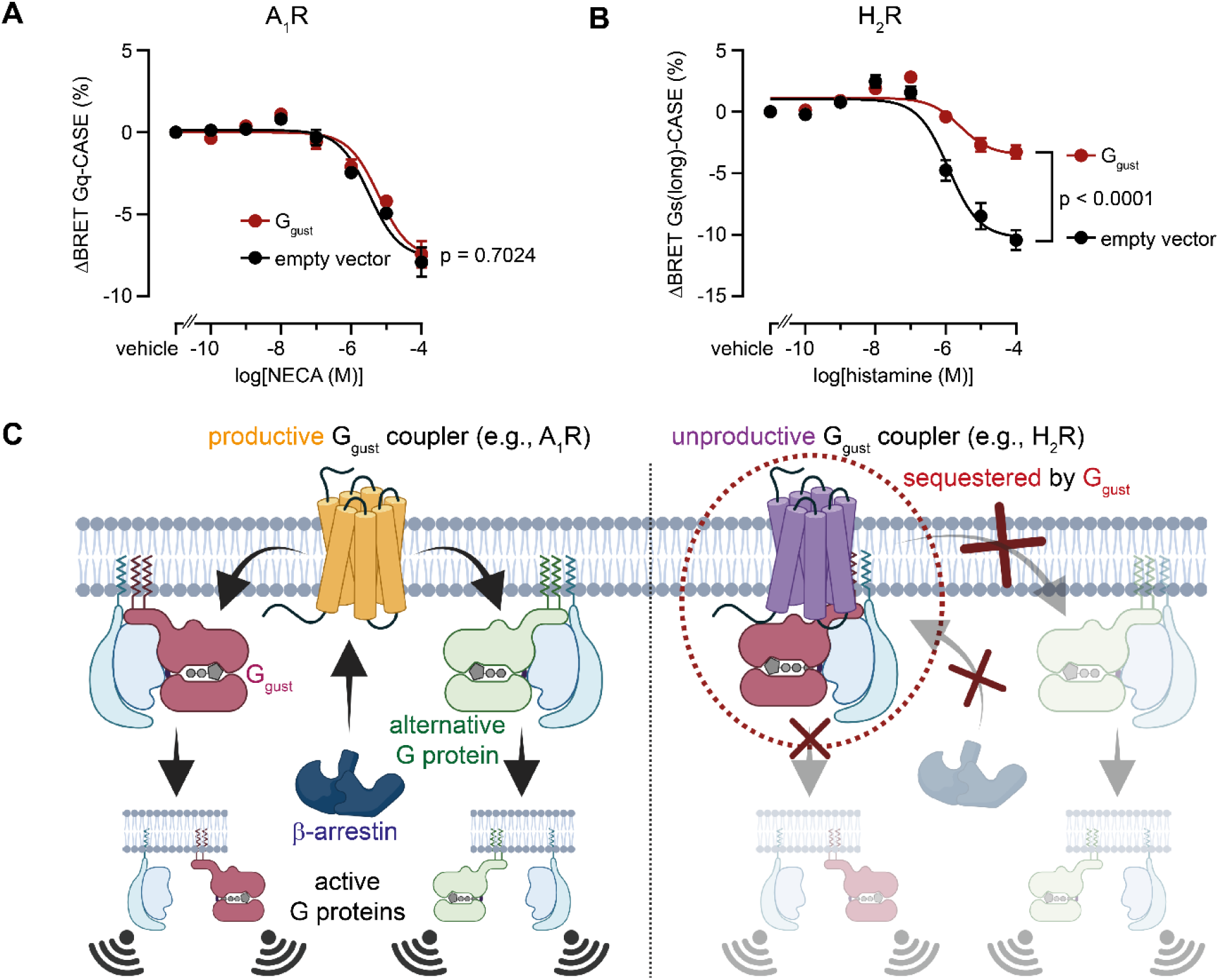
Unproductive GPCR-Ggust complexes suppress activation of alternative G proteins. **A, B)** Concentration-dependent BRET changes induced by NECA (A) and histamine (B) in HEK-293A cells that have been transiently transfected with the A_1_R and Gq-CASE or H_2_R and Gs(long)-CASE, respectively, and empty vector (pcDNA3.1(+)) or G_gust_ (double dark Ggust-CASE). **C)** Scheme of the proposed mechanism leading to the suppression of alternative signaling pathways by unproductive GPCR-G_gust_ complexes. Data show mean ± SEM of three (A) or six (B) independent experiments. Statistical significance was tested using Student’s unpaired t-test of the 100 μM NECA-(A) or 100 μM histamine-induced BRET responses (B).

Overall, these findings, together with our previous data, suggest that unproductive GPCR-G_gust_ complexes sequester receptors away from alternative G proteins, thereby attenuating their activation. Consequently, the reduced release of Gβγ subunits limits the recruitment of cytosolic GRKs to activated receptors[42,43], resulting in diminished β-arrestin engagement with phosphorylated receptors (**Fig. 7C**).

## Discussion

Many GPCRs are able to couple to and activate more than just one type of G proteins in a given cellular setting[28]. While the preferences of GPCRs for certain G protein subtypes have been a body of intense research in past decades, the possibility that different GPCRs can also tune G protein activity into different directions has, however, only recently gained attention with reports of unproductive GPCR-G protein engagements[37,38]. The present study shows that G_gust_ can couple to and be regulated by several non-tastant GPCRs when expressed in HEK-293 cells. Importantly, several receptors reduced basal G_gust_ activity and signal transduction. This observation complements recent reports of unproductive GPCR–G protein complexes and raises intriguing question regarding the physiological consequences of these assemblies in the context of physiology and pathophysiology.

The G protein G_gust_ has long been exclusively viewed as a sensor of taste due to its coupling to tastant-binding GPCRs in gustatory cells. G_gust_’s roles in other parts of the human body remain poorly characterized. Our study begins to illuminate these poorly understood roles and suggests a dual mode of GPCR regulation in cells in which G_gust_ acts as a determinant of divergent signaling outcomes. We show that G_gust_ not only couples to non-tastant GPCRs, but that its interaction with GPCRs can be either productive – leading to enhanced signal transduction – or unproductive, which reduces basal G_gust_-dependent signaling and suppresses the activation of alternative signaling pathways. Previous co-expression analyses suggested that G_gust_ may couple to non-tastant GPCRs in pancreatic β-cells and the colon to promote central physiological responses, such as GLP1 release and insulin secretion, respectively[44,45], but direct evidence for these interactions has been lacking until today.

Our findings raise the question of how suppression of G_gust_ signaling can occur mechanistically and what physiological consequences such inhibition of basal G_gust_ signaling and other pathways may have. Reduction of basal G_gust_ activity and signal transduction requires a sufficient level of basal G_gust_ signaling capacity in cells, as shown for other members of the G_i/o_ family which all elicit high levels of GDP/GTP exchange without external stimulation in biochemical experiments[46,47].

In fact, basal signaling activity of Gα_gust_ is required to maintain low levels of cAMP and insulin secretion in and from pancreatic β-cells[48] and ensures low Ca^2+^ signaling in taste-buds[49], indicating that basal G_gust_ activity is of vital physiological importance across tissues. Productive and unproductive GPCRs coupling to G_gust_ could modulate the sensitivity of these cells prior to additional external (GPCR) stimuli. Likewise, G_gust_-mediated sequestration of GPCRs may serve as a physiological mechanism to prevent concurrent activation of parallel signaling pathways within the same cell in the presence of multiple agonists. Yet, further investigations in more native systems are required to understand the physiological importance of these complexes. In particular, the molecular stoichiometry of GPCRs and different Gα subtypes present in the same cell may influence whether G_gust_ couples to these GPCRs in a physiological setting. Because our experiments were performed in an overexpression system, the relative abundance of receptors and G proteins likely differs from that in native tissues and may therefore affect the formation of these complexes.

On a mechanistic level, it needs to be determined which molecular or atomistic components define a GPCR as a productive or unproductive interaction partner for G_gust_. In our profiling of 24 different GPCRs, we did not observe striking correlations between productive/unproductive G_gust_ coupling and primary coupling preferences across the four G protein families (**Fig. 3**), suggesting that other determinants than those defining G protein coupling preference underlie the productivity of a given GPCR-G_gust_ interaction. While unproductive coupling of GPCRs to G_12_ is influenced by the C-terminal α5 helix of Gα_12_[37], the observation that G_i1_ and G_o1_ – in contrast to G_gust_ – form productive complexes with H_2_R despite the high sequence similarity of the Gα subunits in their C-terminal region (**Fig. S9**), suggests that also other regions influence the signaling capacity of G_gust_ after receptor engagement. Structural studies and mutational mapping of receptor-G_gust_ pairs could provide these mechanistic insights and open avenues to target these interactions.

Together, our work identifies the G protein G_gust_ as a broadly interacting G_i/o_-family member whose signaling output can be either enhanced or suppressed by non-tastant GPCRs, and which can function as a delimiter of alternative signaling pathways. Future studies combining mutational analysis, structural approaches, and experiments in native cell systems will be required to determine whether the regulatory principles uncovered here contribute to physiological processes, such as nutrient sensing and hormone secretion, *in vivo*.

## Material and Methods

### Reagents

Apyrase, grade VI from potato, digitonin and PEI (polyethyleneimine) were obtained from Sigma Aldrich. GPCR agonists were purchased from MedChemExpress or Sigma Aldrich. Furimazine and the Nano-Glo® HiBiT Extracellular Detection System were from Promega. GDPβS and GTPγS were from Jena Bioscience. Cell culture media (DMEM, HBSS, DPBS, FBS, penicillin/streptomycin, L-glutamine, Trypsin-EDTA and OptiMEM) were from Capricorn. All cloning reagents, except for oligonucleotides, were from NEB. Oligonucleotides were obtained from IDT. Cell culture flasks and white 96 well plates were from Greiner Bio-One.

### Plasmids and molecular cloning

Plasmids encoding Gα_gust_-Nluc, Gα_11_-Nluc, Gα_o2_-Nluc, PDE6γ-cpVenus, p63RhoGEF-cpVenus, PDZ-RhoGEF-cpVenus and N-terminally HiBiT-labeled GPCRs were synthesized and subcloned into a pcDNA3.1(+) by Genscript. YFP-labeled Gα effector constructs were kindly provided by Moritz Bünemann (Marburg, Germany). Plasmids encoding β-arrestin2-RlucII and rGFP-CAAX were kindly provided by Michel Bouvier (Montreal, Canada). Ggust-CASE, G11-CASE and Go2-CASE were cloned by sub-cloning the respective Gα-Nluc sequences into the G_o1_-CASE template using Gibson assembly, thereby replacing Gα_o1_-Nluc. The point mutations in cpVenus and Nluc to generate dark Ggust-CASE and double dark Ggust-CASE were introduced into Ggust-CASE using site-directed mutagenesis. Gα_q_-Nluc was PCR-amplified from Gq-CASE[18] and sub-cloned into pcDNA3.1(+) using Gibson assembly. Unlabeled Gα_gust_ was generated through blunt-ended ligation using Gα_gust_-Nluc in pcDNA3.1(+) as a template. Plasmid encoding Venus(NT)-Gβ_1_-Venus(CT)-Gγ_2_ was cloned by replacing the Gγ sequence fragment in Venus(NT)-Gβ_1_-Venus(CT)-Gγ_4_ (synthesized and sub-cloned into pcDNA3.1 (+) by Twist Bioscience) using Gibson assembly. C-terminally Nluc-tagged GPCRs were cloned through PCR amplification of the HiBiT-GPCR fragments and insertion into an Nluc sequence-containing pcDNA3.1(+) template. All plasmids were confirmed by sequencing.

### Cell culture

HEK-293A cells were from ThemoFisher (cat# R70507) and Gα knockout cells were kindly provided by Asuka Inoue[41] (Tohoku University, Miyagi, Japan; Kyoto University, Kyoto, Japan). All cell lines were grown in Dulbecco’s modified Eagle’s medium (DMEM) supplemented with 2 mM glutamine, 10 % fetal calf serum, streptomycin (0.1 mg/mL), and penicillin (100 U/mL) at 37 °C with 5 % CO_2_. Absence of mycoplasma contamination was routinely confirmed by PCR.

### Transient transfection and plating

Resuspended cells (300,000 cells/mL) were transfected in suspension with a total of 1 μg DNA/mL suspension using PEI (1 mg/ml stock solution; 3 μL PEI solution per μg DNA). The exact DNA ratios for co-expression of different proteins are stated in the respective methods sections. Cells mixed with the transfection reagents were seeded onto 96-well plates (100 mL cell suspension / well) and grown for 48 h at 37 °C with 5 % CO_2_. White plates were used for our BRET-based experiments.

### Permeabilization procedure

Cells grown in 96-well microtiter plates were washed with 120 mL HBSS and then incubated with 0.02 U/mL apyrase and 0.001% digitonin (w/v) dissolved in internal buffer (100 mM K^+^-aspartate, 30 mM KCl, 10 mM HEPES, 5 mM EGTA, 1 mM MgCl_2_, 10 mM NaCl, pH 7.35) for 25 minutes at room temperature. Prior to the experiments, permeabilized cells were then supplemented with 1 μM GDPβS and 1/1000 dilution of furimazine (only for BRET experiments).

Luminescence spectra recording of Ggust-CASE and dark Ggust-CASE in intact cells HEK293A cells were transfected with plasmid encoding Ggust-CASE or dark Ggust-CASE. 48 hours post-transfection, cells were washed with 120 μL/well HBSS and incubated with 80 mL/well of a 1/1000 dilution of furimazine for 2 minutes at room temperature. Next, luminescence emission intensities were scanned with 1 nm resolution from 400 nm – 600 nm. All experiments were performed using a ClarioStar Plus plate reader with an integration time of 0.3 seconds.

### Luminescence-based quantification of HiBiT-H_2_R surface expression

HEK293A cells were transfected with 25% HiBiT-H_2_R plasmid, 25% pcDNA and 50% pcDNA or double dark Ggust-CASE. 48 hours post-transfection, intact cells were washed with 120 mL/well HBSS and incubated with a 1/200 dilution of the Nano-Glo® HiBiT Extracellular Substrate and a 1/400 dilution of the LgBiT protein for 30 minutes at room temperature. Nluc emission was then quantified using a ClarioStar Plus plate reader with the following settings: λ(emission): 450-80 nm; integration time: 0.9 seconds.

### BRET measurements in intact cells

HEK293A cells were transfected with 50% receptor plasmid and 50% Ggust-CASE, or a mix of 10% Gα_gust_-Nluc or dark Ggust-CASE with 40% PDE6γ-cpVenus. For competition experiments, 25% receptor plasmid was mixed with 25% G-CASE (Gq-CASE for A_1_R and Gs(long)-CASE for H_2_R) or 5% β-arrestin2-RlucII and 20% rGFP-CAAX[50], along with 50% double dark Ggust-CASE or pcDNA. 48 hours post-transfection, intact cells were washed with 120 mL/well HBSS and incubated with 72 μL/well of a 1/1000 dilution of furimazine for 2 minutes at room temperature. Next, BRET was recorded in three consecutive reads before (serial dilutions of) agonists or vehicle control were added, followed by additional BRET reads. The peak response of the time course data was considered for the generation of concentration response curves. All experiments were performed using a ClarioStar Plus or PheraStar FSX plate reader. For experiments with the BRET donor Nluc and YFP or cpVenus as BRET acceptors, the following settings were used: λ(donor emission): 450-80 nm; λ(acceptor emission): 535-30 nm; integration time: 0.3 seconds. For experiments with the BRET donor RlucII and GFP as BRET acceptor, the following settings were used: λ(donor emission): 400-100 nm; λ(acceptor emission): 520-30 nm; integration time: 0.9 seconds.

### BRET measurements in permeabilized cells

HEK293A cells were transfected with 50% pcDNA and Ggust-CASE, or 10% dark Ggust-CASE with 40% PDE6γ-cpVenus. 48 hours post-transfection, cells were permeabilized and nucleotide depleted by incubating with 0.001 % (m/v) digitonin and 0.02 U/ml apyrase for 25 minutes at room temperature. Then, permeabilized cells were loaded with 1 μM GDPβS and 1/1000 furimazine. Next, BRET was recorded in several consecutive reads before 10 μM GTPγS or vehicle control were added with automated injectors, followed by additional BRET reads. All experiments were performed using a PheraStar FSX plate reader with the following settings: λ(donor emission): 450-80 nm; λ(acceptor emission): 535-30 nm; integration time: 0.3 seconds.

### Nucleotide-sensitivity of GPCR-Nluc/G_gust_ complexes in permeabilized cells

HEK293A cells lacking endogenous Gα proteins[41] were transfected with 25% GPCR-Nluc, 25% unlabeled Gα_gust_ and 50% Venus-Gβ_1_γ_2_. 48 hours post-transfection, cells were permeabilized, loaded with 1/1000 furimazine and saturating concentrations of GPCR agonists (i.e., 10 μM bradykinin for BK_2_R; 100 μM NECA for A_1_R and A_2A_R; 100 μM serotonin for 5HT_1A_R; 100 μM histamine for H_2_R and H_3_R). After five minutes of incubation at room temperature, BRET was recorded in several consecutive reads before 10 μM GTPγS or vehicle control were added with automated injectors, followed by additional BRET reads. All experiments were performed using a PheraStar FSX plate reader with the following settings: λ(donor emission): 450-80 nm; λ(acceptor emission): 535-30 nm; integration time: 0.3 seconds.

### Quantification and statistical analysis

BRET was defined as acceptor emission over donor emission. All baseline BRET values were averaged to obtain BRET_basal_. Raw ΔBRET at any given time point t were calculated as follows: raw ΔBRET_t_ = ((BRET_t_-BRET_basal_)/BRET_basal_)*100. The average raw ΔBRET_t_ of all vehicle-treated technical replicates were calculated for any given time point t. These values were subtracted from all raw ΔBRET_t_ to obtain corrected ΔBRET_t_ values (named only “ΔBRET” in figure axes for simplicity) for each test condition. Corrected ΔBRET values were plotted over time and time points with saturating response of all concentrations tested were selected. Semi-logarithmic concentration-response curves were plotted and fitted using a sigmoidal concentration–response model (with the Hill slope fixed to +1 or −1). For the data in Fig. 3, non-G_gust_-couplers (nc) were identified by comparing a linear model with a zero slope to a sigmoidal concentration–response model using an extra sum of squares F test (p > 0.05).

Tissue-level GPCR expression analysis in the small intestine and duodenum analysis We queried the Human Protein Atlas with the following prompts:

1. protein_class:G-protein coupled receptors;Any AND normal_expression:Duodenum;Any
2. protein_class:G-protein coupled receptors;Any AND normal_expression:Small intestine;Any The results were exported as TSV files and imported into Microsoft Excel for inspection (compiled in **Supplementary Data 1**).

### Single-cell-level co-expression analysis of GPCRs with Gα_gust_

Publicly available single cell RNAseq data were obtained from the human protein atlas[15]. Data processing and analysis were performed using R. Read count tables for 36 datasets and the respective metadata were downloaded before co-expression of *GNAT3* and the selected GPCRs was analysed. A gene was considered expressed if the read count was greater than zero. Cells were classified into four categories: both genes expressed, only *GNAT3* expressed, only the GPCR expressed, or neither gene expressed. Cell counts were calculated for each cell subtype and tissue.

## Acknowledgements

The results shown here are in whole or part based upon data generated by the TCGA Research Network: https://www.cancer.gov/tcga. We thank Asuka Inoue (Kyoto University, Tohoku Universiy, Japan) for providing the Gα knockout HEK-293A cells. This project has received funding from the Deutsche Forschungsgemeinschaft (DFG) under the Emmy Noether grant agreement number 542889291 (GZ: SCHI 1508/3-1).

## Data availability

All data generated or analyzed during this study are included in this published article and its supplementary information files. The data that support the findings of this study are available from the corresponding author (H.S.) upon reasonable request.

## Author contributions

V.J.A., M.S.T., S.J.B.G., and H.S. conducted experiments. V.J.A., M.S.T., S.J.B.G., I.K., D.H., and H.S. designed experiments and analyzed data. H.S. wrote the manuscript with input from all authors. H.S. conceived and supervised the project.

## Supplementary Information

Supplementary Data 1 – Expression of GPCRs in the duodenum or small intestine Supplementary Data 2 – Single-cell-level co-expression analysis of GPCRs and Gα_gust_ Supplementary Information with Figures S1 – S7

