## Supporting information for "Functional diversity of GPCR-Gustducin complexes controls signaling output and suppresses alternative pathways"

for

#### Figure S1

**$G\alpha_{gust}$-Nluc**:

MGSGISSESKESAKRSKELEKKLQEDAERDARTVKLLLLGAGESGKSTIVKQMKIIHK
NGYSEQECMEFKAVIYSNTLQSILAIVKAMTTLGIDYVNPRSAEDQRQLYAMANTLED
SGGGGTRSVFTLEDFVGDWRQTAGYNLDQVLEQGGVSSLFQNLGVSVTPIQRIVLS
GENGLKIDIHVIIPYEGLSGDQMGQIEKIFKVVYPVDDHHFKVILHYGTLVIDGVTPNMI
DYFGRPYEGIAVFDGKKITVTGTLWNGNKIIDERLINPDGSLLFRVTINGVTGWRLCE
RILATGGGGSGGMTPQLAEVIKRLWRDPGIQACFERASEYQLNDSAAYYLNDLDRIT
ASGYVPNEQDVLHSRVKTTGIIETQFSFKDLHFRMFDVGGQRSERKKWIHCFEGVT
CIIFCAALSAYDMVLVEDEEVNRMHESLHLFNSICNHKYFSTTSIVLFLNKKDIFQEKV
TKVHLSICFPEYTGPNTFEDAGNYIKNQFLDLNLKKEDKEIYSHMTCATDTQNVKFVF
DAVTDIIIKENLKDCGLF*

**$G\beta_3$-T2A-cpVenus-$G\gamma_9$**:

MGEMEQLRQEAEQLKKQIADARKACADVTLAELVSGLEVVGRVQMRTRRTLRGHL
AKIYAMHWATDSKLLVSASQDGKLIVWDSYTTNKVHAIPLRSSWVMTCAYAPSGNFV
ACGGLDNMCSIYNLKSREGNVKVSRELSAHTGYLSCCRFLDDNNIVTSSGDTTCAL
WDIETGQQKTVFVGHTGDCMSLAVSPDFNLFISGACDASAKLWDVREGTCRQTFT
GHESDINAICFFPNGEAICTGSDDASCRLFDLRADQELICFSHESIICGITSVAFSLSG
RLLFAGYDDFNCNVWDSMKSERVGILSGHDNRVSCLGVTADGMAVATGSWDSFLKI
WNEGRGSLLTCGDVEENPGPTGMDGGVQLADHYQQNTPIGDGPVLLPDNHYLSY
QSALSKDPNEKRDHMVLLEFVTAAGITLGMDELYKGGSGGMVSKGEELFTGVVPILV
ELDGDVNGHKFSVSGEGEGDATYGKLTLKLICTTGKLPVPWPTLVTTLGYGLQCFAR
YPDHMKQHDFFKSAMPEGYVQERTIFFKDDGNYKTRAEVKFEGDTLVNRIELKGIDF
KEDGNILGHKLEYNYNSHNVYITADKQKNGIKANFKIRHNIESGLRSAQDLSEKDLLK
MEVEQLKKEVKNTRIPISKAGKEIKEYVEAQAGNDPFLKGIPEDKNPFKEKGGCLIS*

**Figure S1. Amino acid sequence of $G_{gust}$-CASE.**

**Fig. S2**

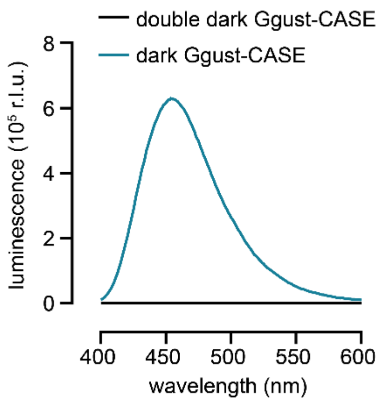


**Figure S2. Luminescence emission spectra of dark Ggust-CASE and double dark Ggust-CASE.** Data represents the emission intensity in one replicate conducted in HEK-293A cells transiently transfected with plasmids encoding the indicated proteins.

#### Figure S3

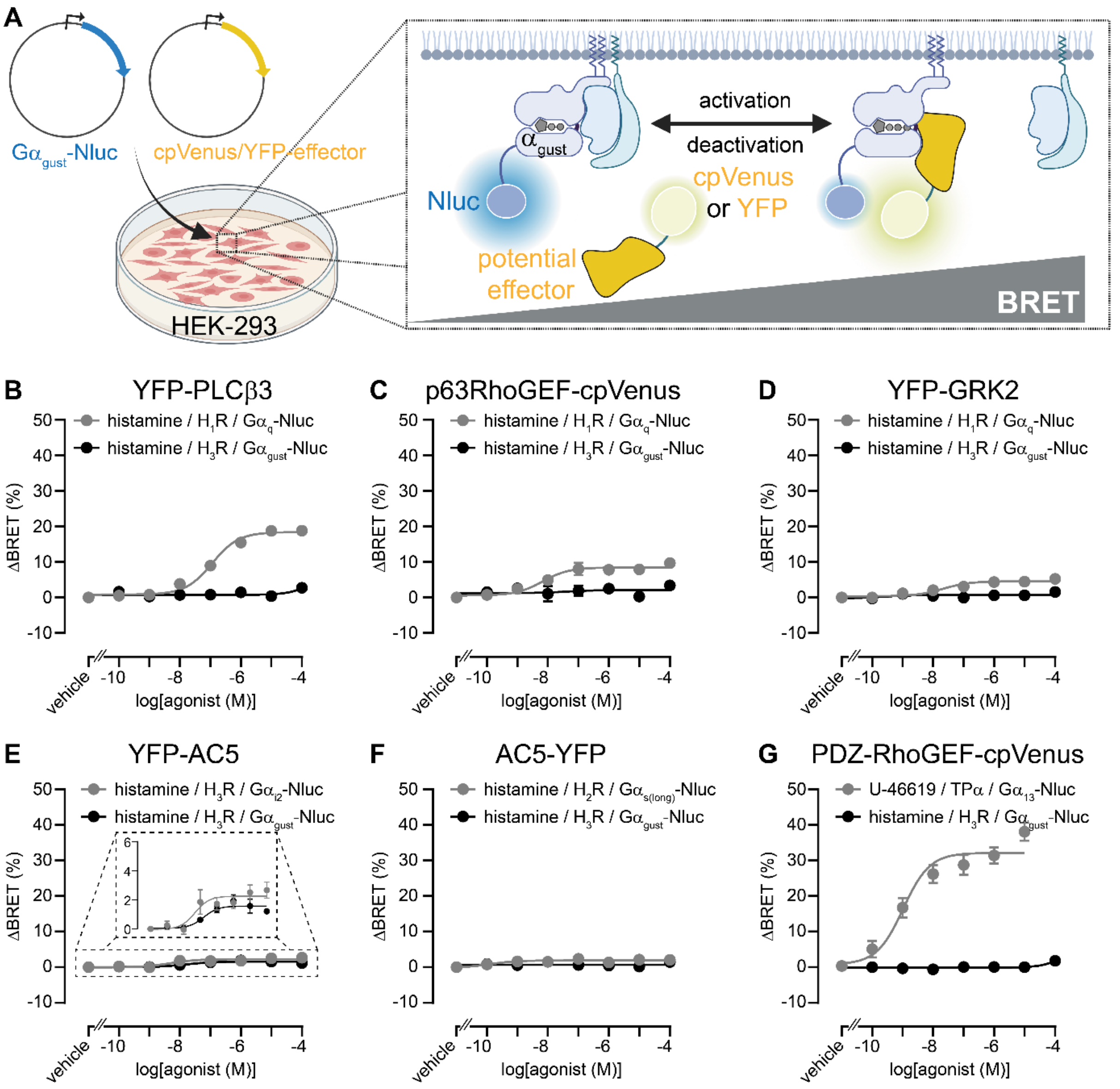


**Figure S3. Testing Gα effector proteins for interactions with Gα$_{gust}$. A)** Scheme of the BRET-based protein-protein interaction assay. **B-G)** Concentration-dependent BRET changes induced by GPCR agonists. Data represent mean ± SEM of three to five biological replicates conducted in HEK-293A cells transiently transfected with plasmids encoding the indicated proteins.

#### Figure S4

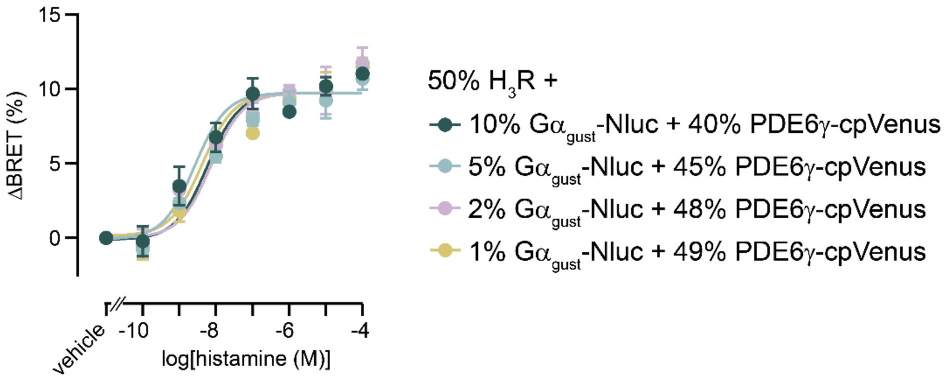


**Figure S4. BRET donor/acceptor titration for the $G\alpha_{gust}$-PDE6γ interaction assay.** Histamine-induced, concentration-dependent BRET changes in HEK-293A cells transiently transfected with different plasmid ratios of $H_3R$, $G\alpha_{gust}$-Nluc and PDE6γ-cpVenus. Data show mean ± SEM of three independent experiments.

**Fig. S5**

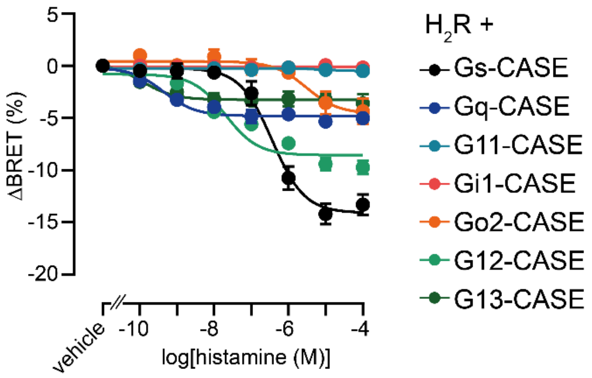


**Figure S5. $H_2R$-mediated G-CASE responses.** Histamine-induced, concentration-dependent BRET changes in HEK-293A cells transiently transfected with the $H_2R$ and different G protein biosensors. Data show mean ± SEM of three independent experiments.

#### Figure S6

### A

**$G\alpha_{11}$-Nluc**:

MTLESMMACCLSDEVKESKRINAEIEKQLRRDKRDARRELKLLLLGTGESGKSTFIK
QMRIIHGAGYSEEDKRGFTKLVYQNIFTAMQAMIRAMETLKILYKYEQNKANALLIRE
VDVEKVTTFEHQYVSAIKTLWEDPGIQECYDRRREYQLSDSAKYYLTDVDRIATLGY
LPTQQDVLRVRVPTTGIIEYPFDLENIIFRMVDVGGQRSERRKWIHCFENVTSIMFLV
ALSEYDQVLVESDNESGGGGTRSVFTLEDFVGDWRQTAGYNLDQVLEQGGVSSLF
QNLGVSVTPIQRIVLSGENGLKIDIHVIIPYEGLSGDQMGQIEKIFKVVYPVDDHHFKVI
LHYGTLVIDGVTPNMIDYFGRPYEGIAVFDGKKITVTGTLWNGNKIIDERLINPDGSLL
FRVTINGVTGWRLCERILATGGGGSNRMEESKALFRTIITYPWFQNSSVILFLNKKDL
LEDKILYSHLVDYFPEFDGPQRDAQAAREFILKMFVDLNPDSDKIIYSHFTCATDTENI
RFVFAAVKDTILQLNLKEYNLV*

**$G\beta_3$-T2A-cpVenus-$G\gamma_9$**:

MGEMEQLRQEAEQLKKQIADARKACADVTLAELVSGLEVVGRVQMRTRRTLRGHL
AKIYAMHWATDSKLLVSASQDGKLIVWDSYTTNKVHAIPLRSSWVMTCAYAPSGNFV
ACGGLDNMCSIYNLKSREGNVKVSRELSAHTGYLSCCRFLDDNNIVTSSGDTTCAL
WDIETGQQKTVFVGHTGDCMSLAVSPDFNLFISGACDASAKLWDVREGTCRQTFT
GHESDINAICFFPNGEAICTGSDDASCRLFDLRADQELICFSHESIICGITSVAFSLSG
RLLFAGYDDFNCNVWDSMKSERVGILSGHDNRVSCLGVTADGMAVATGSWDSFLKI
WNEGRGSLLTCGDVEENPGPTGMDGGVQLADHYQQNTPIGDGPVLLPDNHYLSY
QSALSKDPNEKRDHMVLLEFVTAAGITLGMDELYKGGSGGMVSKGEELFTGVVPILV
ELDGDVNGHKFSVSGEGEGDATYGKLTLKLICTTGKLPVPWPTLVTTLGYGLQCFAR
YPDHMKQHDFFKSAMPEGYVQERTIFFKDDGNYKTRAEVKFEGDTLVNRIELKGIDF
KEDGNILGHKLEYNYNSHNVYITADKQKNGIKANFKIRHNIESGLRSAQDLSEKDLLK
MEVEQLKKEVKNTRIPISKAGKEIKEYVEAQAGNDPFLKGIPEDKNPFKEKGGCLIS*

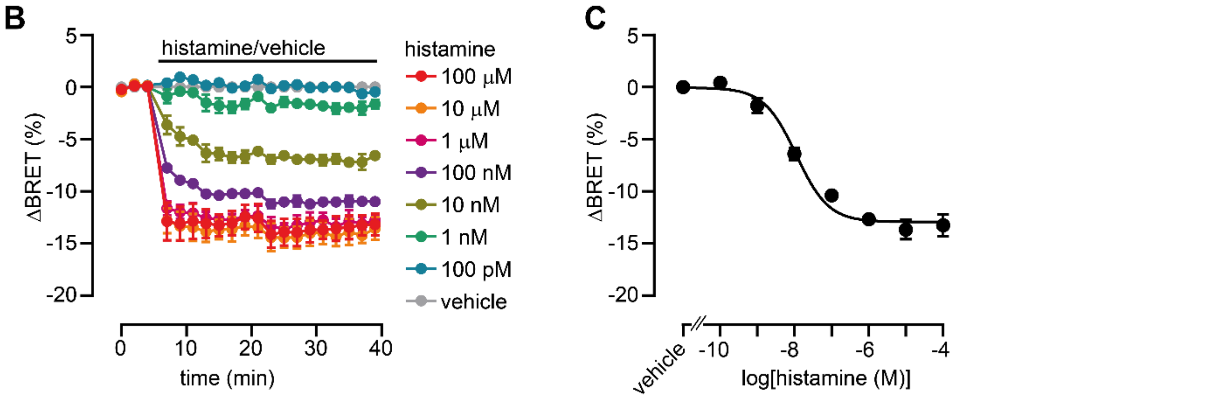


**Figure S6. Design and validation of G11-CASE. A)** Amino acid sequence of G11-CASE. **B, C)** Time- and concentration-dependent BRET changes induced by histamine in HEK-293A cells that have been transiently transfected with the $H_1R$ and G11-CASE. Data show mean ± SEM of three independent experiments.

**Figure S7**

**$G\alpha_{o2}$-Nluc**:

MGCTLSAEERAALERSKAIEKNLKEDGISAAKDVKLLLLGAGESGKSTIVKQMKIIHE
DGFSGEDVKQYKPVVYSNTIQSLAAIVRAMDTLSGGGGTRSVFTLEDFVGDWRQTA
GYNLDQVLEQGGVSSLFQNLGVSVTPIQRIVLSGENGLKIDIHVIIPYEGLSGDQMGQ
IEKIFKVVYPVDDHHFKVILHYGTLVIDGVTPNMIDYFGRPYEGIAVFDGKKITVTGTL
WNGNKIIDERLINPDGSLLFRVTINGVTGWRLCERILATGGGGSGIEYGDKERKADA
KMVCDVVSRMEDTEPFSAELLSAMMRLWGDSGIQECFNRSREYQLNDSAKYYLDS
LDRIGAADYQPTEQDILRTRVKTTGIVETHFTFKNLHFRLFDVGGQRSERKKWIHCF
EDVTAIIFCVALSGYDQVLHEDETTNRMHESLKLFDSICNNKWFTDTSIILFLNKKDIF
EEKIKKSPLTICFPEYTGPSAFTEAVAYIQAQYESKNKSAHKEIYTHVTCATDTNNIQF
VFDAVTDVIIAKNLRGCGLY*

**$G\beta_3$-T2A-cpVenus-$G\gamma_9$**:

MGEMEQLRQEAEQLKKQIADARKACADVTLAELVSGLEVVGRVQMRTRRTLRGHL
AKIYAMHWATDSKLLVSASQDGKLIVWDSYTTNKVHAIPLRSSWVMTCAYAPSGNFV
ACGGLDNMCSIYNLKSREGNVKVSRELSAHTGYLSCCRFLDDNNIVTSSGDTTCAL
WDIETGQQKTVFVGHTGDCMSLAVSPDFNLFISGACDASAKLWDVREGTCRQTFT
GHESDINAICFFPNGEAICTGSDDASCRLFDLRADQELICFSHESIICGITSVAFSLSG
RLLFAGYDDFNCNVWDSMKSERVGILSGHDNRVSCLGVTADGMAVATGSWDSFLKI
WNEGRGSLLTCGDVEENPGPTGMDGGVQLADHYQQNTPIGDGPVLLPDNHYLSY
QSALSKDPNEKRDHMVLLEFVTAAGITLGMDELYKGGSGGMVSKGEELFTGVVPILV
ELDGDVNGHKFSVSGEGEGDATYGKLTLKLICTTGKLPVPWPTLVTTLGYGLQCFAR
YPDHMKQHDFFKSAMPEGYVQERTIFFKDDGNYKTRAEVKFEGDTLVNRIELKGIDF
KEDGNILGHKLEYNYNSHNVYITADKQKNGIKANFKIRHNIESGLRSAQDLSEKDLLK
MEVEQLKKEVKNTRIPISKAGKEIKEYVEAQAGNDPFLKGIPEDKNPFKEKGGCLIS*

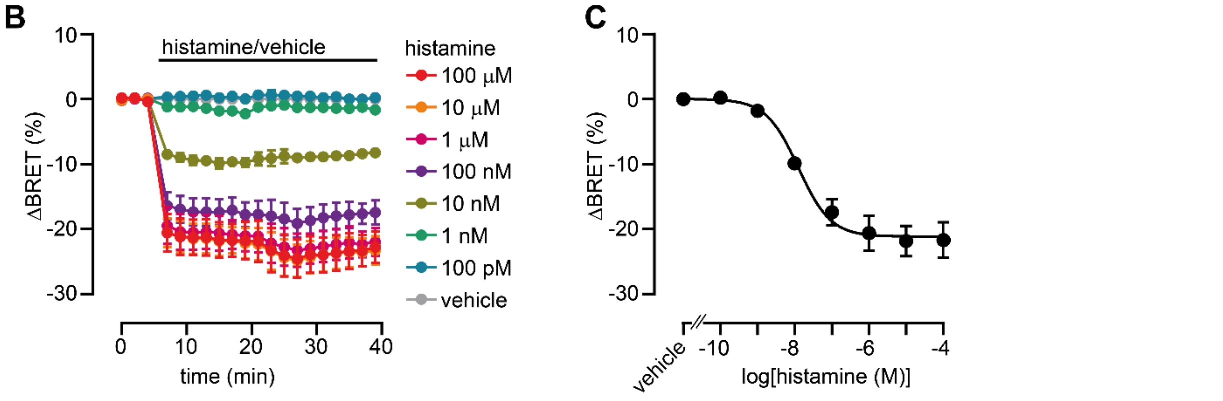


**Figure S7. Design and validation of Go2-CASE. A)** Amino acid sequence of Go2-CASE. **B, C)** Time- and concentration-dependent BRET changes induced by histamine in HEK-293A cells that have been transiently transfected with the $H_3R$ and Go2-CASE. Data show mean ± SEM of three independent experiments.

#### Fig. S8

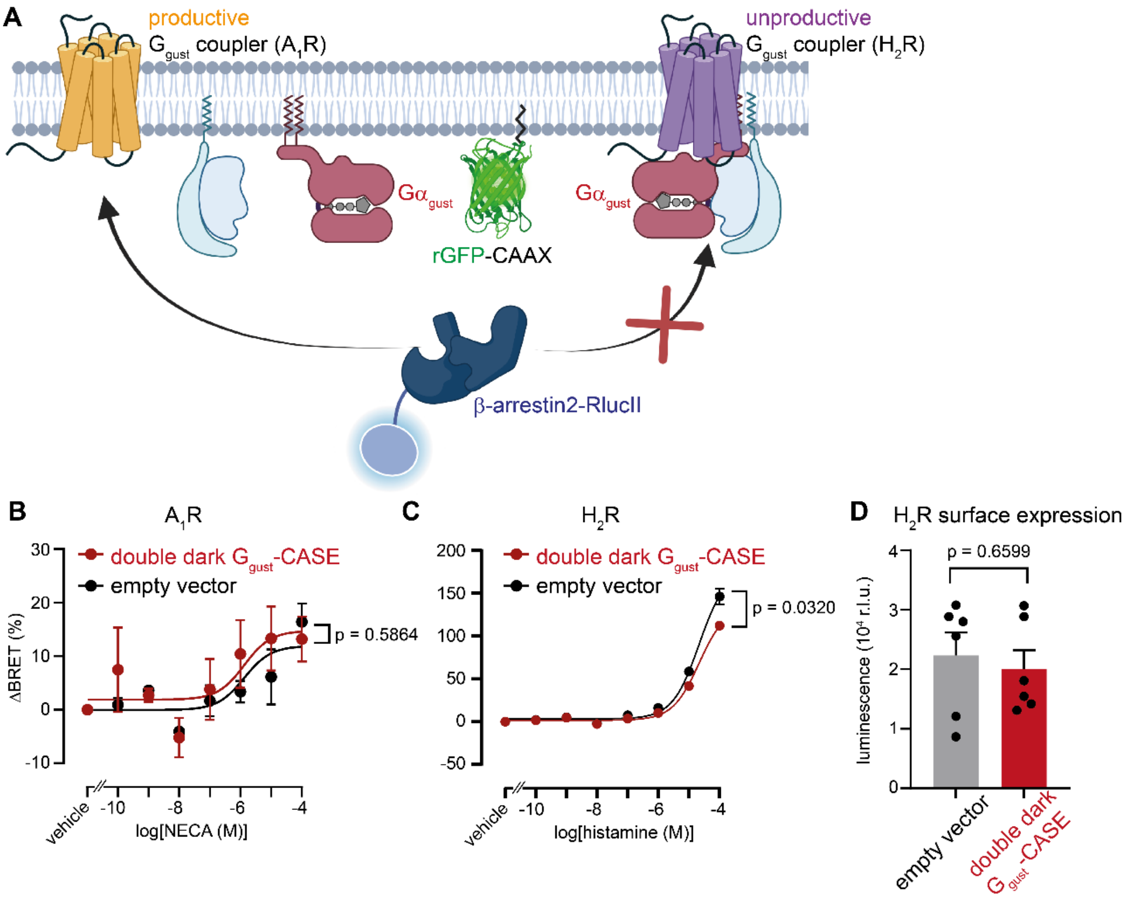


**Figure S8. Unproductive GPCR-Ggust complexes suppress β-arrestin2 recruitment. A)** Scheme of the intermolecular BRET assay. **B, C)** Concentration-dependent BRET changes induced by NECA (B) and histamine (C) in HEK-293A cells that have been transiently transfected with the $A_1R$ or $H_2R$, respectively, and the plasmids encoding rGFP-CAAX, β-arrestin2-RlucII and empty vector (pcDNA3.1(+)) or double dark Ggust-CASE. **D)** Luminescence intensity upon HiBiT-LgBiT complementation representing the surface expression levels of $H_2R$ in HEK-293A cells transiently transfected with the HiBiT-$H_2R$ and empty vector (pcDNA3.1(+)) or double dark Ggust-CASE. Data show mean ± SEM of three (B, C) or six (D) independent experiments. Statistical significance was tested using Student's unpaired t-test (in B and C of the 100 μM NECA- or 100 μM histamine-induced responses, respectively).

### Fig. S9

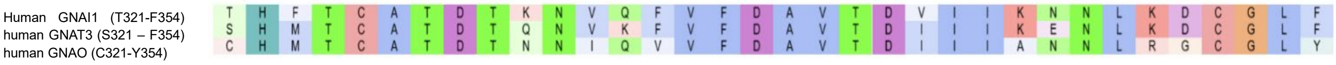


**Figure S9. Sequence alignment of the c-terminal ends of G$\alpha$ subunits G$\alpha_{i1}$, G$\alpha_{gust}$ and G$\alpha_{o1/2}$.**
